# Morphology and immunohistochemical characteristics of the otic ganglion in the chinchilla (*Chinchilla laniger* Molina)

**DOI:** 10.1101/2020.11.12.379503

**Authors:** Waldemar Sienkiewicz, Jacek Kuchinka, Agnieszka Dudek, Elżbieta Nowak, Jerzy Kaleczyc, Małgorzata Radzimirska, Aleksander Szczurkowski

**Author notes:** Corresponding author: W. Sienkiewicz.

## Abstract

The available literature provides relatively little information on the morphology of the autonomic head ganglia in rodents including their neurochemical codding. The present study was thus designed to investigate the morphology and neurochemical properties of the otic ganglion in the chinchilla. The results will contribute to our knowledge of the organization of the autonomic nerve system in mammals.

Morphological investigations of the otic ganglion were performed using the modified acetylcholinesterase method. The cellular structure was investigated with histological techniques and neurochemical properties were studied with double-labelling immunofluorescence method

Macromorphological investigations allowed the otic ganglion to be identified as a compact, oval agglomeration of neurons and nerve fibers located inside the skull on the medial surface of the mandibular nerve, just above the oval foramen. Multidimensional cross-sections revealed densely arranged neuronal perikarya and two populations of nerve cells differing in size were distinguished. The large cells (40–50 μm) accounted for about 80% of the neurons in the otic ganglion cross-sections. Moreover, a small number of intraganglionic nerve fibers was observed. Immunohistochemical staining revealed that over 85% of the neuronal cell bodies in the otic ganglion contained immunoreactivity to VAChT or ChAT. VIP-immunoreactive perikarya comprised approximately 10% of the ganglionic cells. Double staining revealed the presence of VAChT and NOS-positive neurons which amounted to about 45% of the nerve cells in the otic ganglion. NOS-positive only perikarya comprised approx. 15% of all the neurons. Immunoreactivity to enkephalin, substance P, somatostatin and galanin was expressed in single nerve cell bodies and nerve fibres except numerous SP-positive intraganglionic nerve fibres. Some of them stained also for CGRP. Single neurons stained for TH.

The present results, compared with previous findings, suggest the existence interspecies differences in the morphology, cellular structure and immunohistochemical properties of the head autonomic ganglia in mammals.

## Introduction

The use of a modified classical histochemical method to localize cholinesterase activity has allowed autonomic cranial ganglia, including the otic ganglion, to be identified in many species of mammals [1, 2]. Acetylcholinesterase (AChE) positive neurons were found in the otic ganglion in a number of animal species, such as dog [3], cat [4], spotted suslik [5], Egyptian spiny mouse [6], and rat [7]. Comparative anatomical studies performed in mammals indicate the presence of a significant relationship between the morphology, topography, and cellular structure of the parasympathetic head ganglia and the systematic position of the species investigated. Moreover, these ganglia display pronounced immunohistochemical differentiation between species. The specific immunohistochemical markers of parasympathetic structures are the enzyme that synthesizes acetylcholine, choline acetyltransferase (ChAT), and the specific neurotransmitter transporter protein, vesicular acetylcholine transporter (VAChT), responsible for loading acetylcholine into secretory vesicles in neurons, making it available for secretion. The presence of these two substances has been demonstrated in the otic ganglion of the rat [8–11] and pig [12]. Moreover, the expression of other biologically active substances, such as neuropeptide Y, vasoactive intestinal polypeptide, substance P, and calcitonin gene-related peptide, has been found in the otic ganglion of humans, pigs, and rats [13, 8, 12, 9, 10].

The available literature contains no information on the morphology and neurochemical properties of the otic ganglion in the chinchilla. The present study was thus designed to investigate this. The results will contribute to our knowledge of the organization of the autonomic nerve system in mammals.

## Material and methods

The investigations were carried out on twelve chinchilla of both sexes (*Chinchilla laniger* Molina). These studies were approved in accordance with the appropriate Polish statute law on protection of research animals (*Ustawa z dnia 15 stycznia 2015 r. o ochronie zwierząt* …); studies on tissues obtained post-mortem do not require an approval of the Ethics Committee. The material was collected immediately after industrial slaughter at a chinchilla fur farm. The mandibular nerves were exposed under a stereomicroscope, rinsed in physiological solution, and fixed for 30 min. in 10% formalin. Four individuals were used for morphological investigations using the modified acetylcholinesterase method [1, 2]. Another four animals were used for the histological study. Once fixed in formalin, the mandibular nerves were embedded in paraffin (Paraplast plus) and in Historesine® (70-2218-501 Mounting Medicum), cut with a microtome (Mikrotom 335) for 3–5 μm slides, stained according to methylene blue and the H&E methods, and silvered method. Morphometry and photographic documentation were performed using a Nikon Digital Sight SD-L1 System and Nis-Elements 3.22 software with Nikon Eclipse 90i microscope.

The final four animals used for the immunohistochemical investigations were transcardially perfused with 0.4 l of 4% ice-cold buffered paraformaldehyde, and the mandibular nerves were collected as described above. The tissues were postfixed by immersion in the same fixative for 2 hours, rinsed with phosphate buffer and transferred to 30% buffered sucrose solution for storage until further processing. The ganglia were cut into 12 μm - thick cryostat sections, which were processed for double-labelling immunofluorescence method on slide-mounted sections. The sections were washed thrice for 10 min. in PB, incubated 45 min. in 10% normal horse serum (NHS, Cappel, Warsaw, Poland) or normal goat serum (NGS, Cappel, Warsaw, Poland) in PBS containing 0.25% Triton X-100 (Sigma, USA), and incubated overnight at room temperature (RT) with antibodies (Table 1) diluted in PB containing 0.25% Triton X-100. After incubation with primary antiserum, the sections were washed three times for 10 min. in PB and further incubated with secondary antisera for 1 h in RT. After incubation, the sections were washed three times for 10 min. in PB, coverslipped with buffered glycerol, and examined under a confocal microscope (Zeiss, LSM 700). Every seventh section was used to avoid double counting of the same neurons and sixty histological sections were used for the measurement.

**Table 1.**
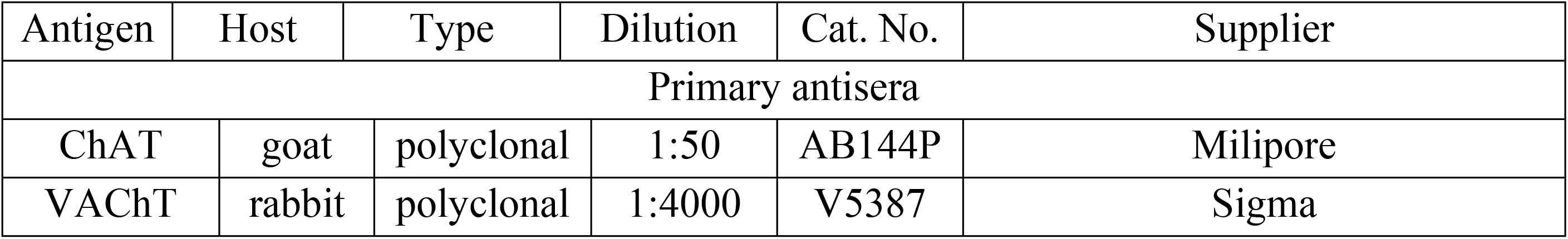
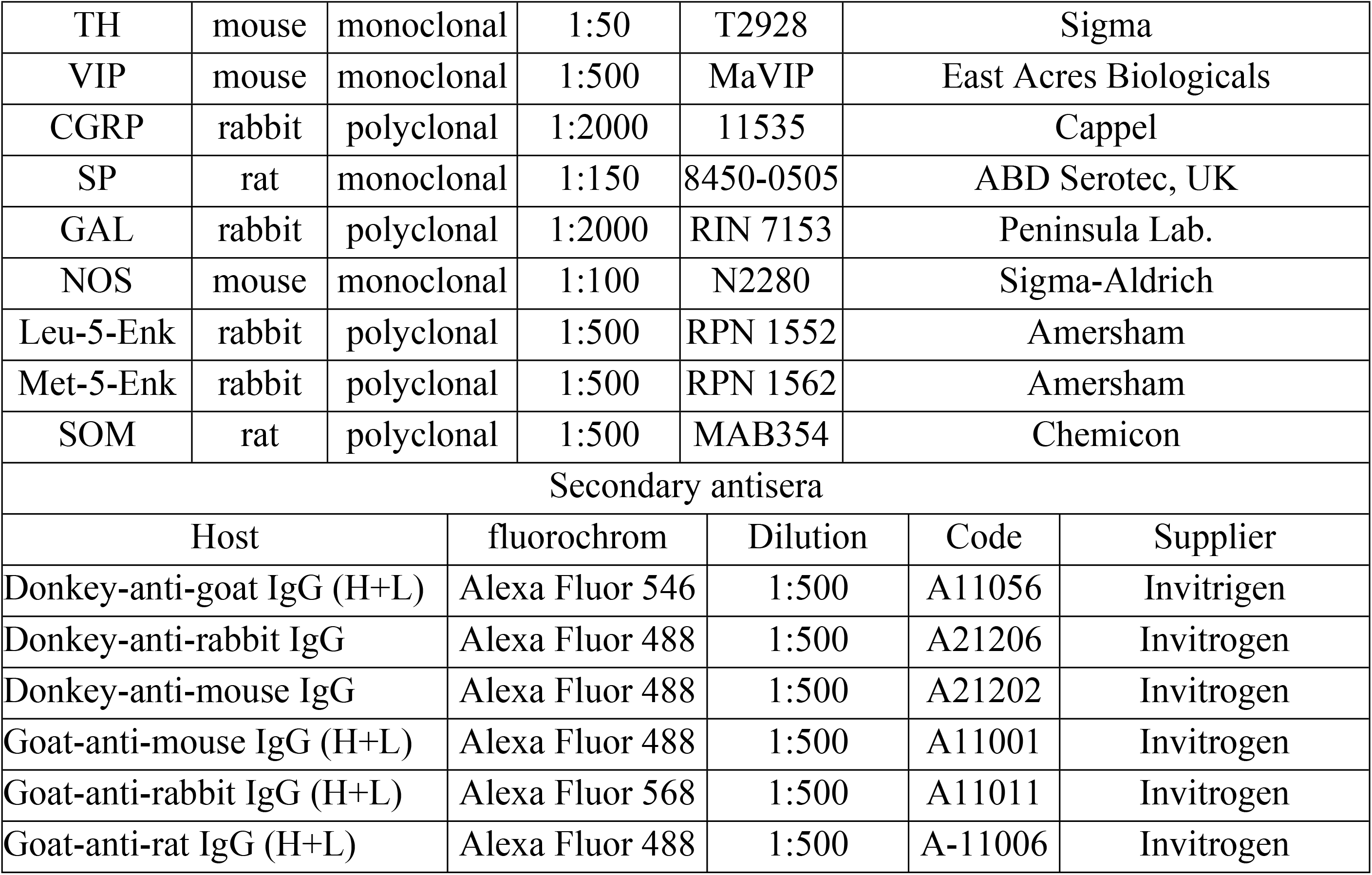
Antisera used in the study.

Staining specificity was controlled by preabsorbtion of a diluted antiserum with 20 μg/ml of an appropriate antigen, which abolished the specific immunoreaction completely. In addition, experiments were carried out in which the primary antiserum was replaced by nonimmune serum or by PBS, in order to verify the specificity of particular immunoreactions.

## Results

The histochemical investigations revealed extensive acetylcholinesterase (AChE) staining in the otic ganglion of the chinchilla. This allowed the ganglion to be identified as a compact, oval agglomeration of neurons and nerve fibers located inside the skull on the medial surface of the mandibular nerve, just above the oval foramen (Fig 1A). The ganglion was 3–5 mm long, 2–3 mm wide and about 0.6 mm thick. The small petrosal nerve, as the parasympathetic radix, reaches the caudal part of the ganglion. In its course, a small aggregation (1 mm long and 0.4 mm wide) of neurocytes was observed. The maxillary artery ran below the ganglion crossing the mandibular nerve more often on the medial side (70% cases), and rarely on the lateral side (Fig 1A).

**Fig 1.**
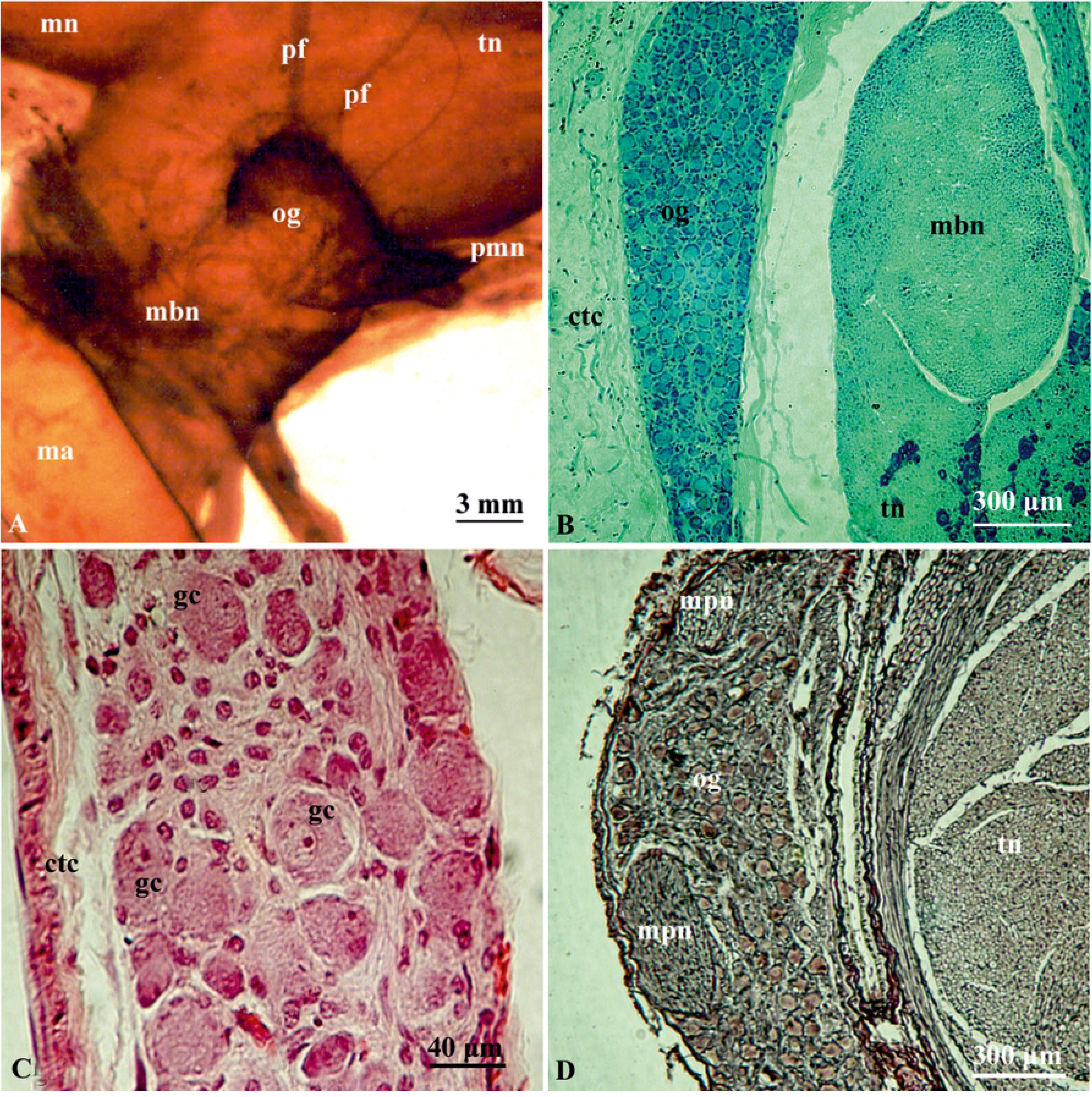
Morphology, topography and cellular structure of the otic ganglion in chinchilla. A) Morphology of the otic ganglion in chinchilla. Thiocholine method. B) Cross-section through the otic ganglion in chinchilla. Methylene blue staining. C) Cross-section through the central part of the otic ganglion in chinchilla. H&E method. D) Cross-section through the otic ganglion in chinchilla. Silver method. og – otic ganglion, pmn – minor petrosal nerve, pf – postganglionic fibres, tn – trigeminal nerve, mn – maxillary nerve, mbn – mandibular nerve, ma – maxillary artery, ctc – connective tissue capsule gc – ganglionic cells, mpn - medial petrosal nerve

Postganglionic fibers leaving the otic ganglion intensely stained for AChE. A delicate bundle of fibers left the superior surface of the structure and entered the trigeminal ganglion (Fig 1A). A small aggregation of neurons in the course of this nerve was frequently observed.

The histological investigations confirmed the close contact between the otic ganglion and the mandibular nerve (Fig 1A-D). Multidimensional cross-sections revealed densely arranged neuronal perikarya. Two populations of nerve cells differing in size could be distinguished: the large neurons with a diameter of 40–50 μm, and the small cells with a diameter of 28–25 μm (Fig 1B, C). The large cells accounted for about 80% of the neurons in the otic ganglion cross-sections. The ganglionic neurons were surrounded by satellite cells with intensely stained nuclei (Fig 1C). Moreover, a small number of intraganglionic nerve fibers was observed (Fig 1D). The entire chinchilla otic ganglion was surrounded by thin connective tissue capsule.

Immunohistochemical staining revealed that over 85% of the neuronal cell bodies in the otic ganglion had immunoreactivity to VAChT (Fig 2A), and a comparable number of the perikarya stained for ChAT (Fig 2C). These cells were mainly of the large type. VIP-immunoreactive (IR) perikarya made up approximately 10% of the ganglionic cells; these were small or medium-sized neurons (Fig 2A). Double staining revealed that approximately 20% of the VIP-IR neurons were VAChT-negative (Fig 2A). Intraganglionic VAChT-IR nerve fibers were numerous and often formed basket-like structures surrounding all the neuronal perikarya (Fig 2A). VIP-positive nerves terminals were scarce. Double staining revealed the presence of VAChT and NOS-positive neurons which amounted to about 45% of the nerve cells in the otic ganglion. NOS-positive only perikarya made up approximately 15% of all the neurons (Fig 2B); these neurons were cells of the small type, whereas double stained neurons were medium and large perikarya. Only few intraganglionic NOS-positive nerve fibers were encountered.

**Fig 2.**
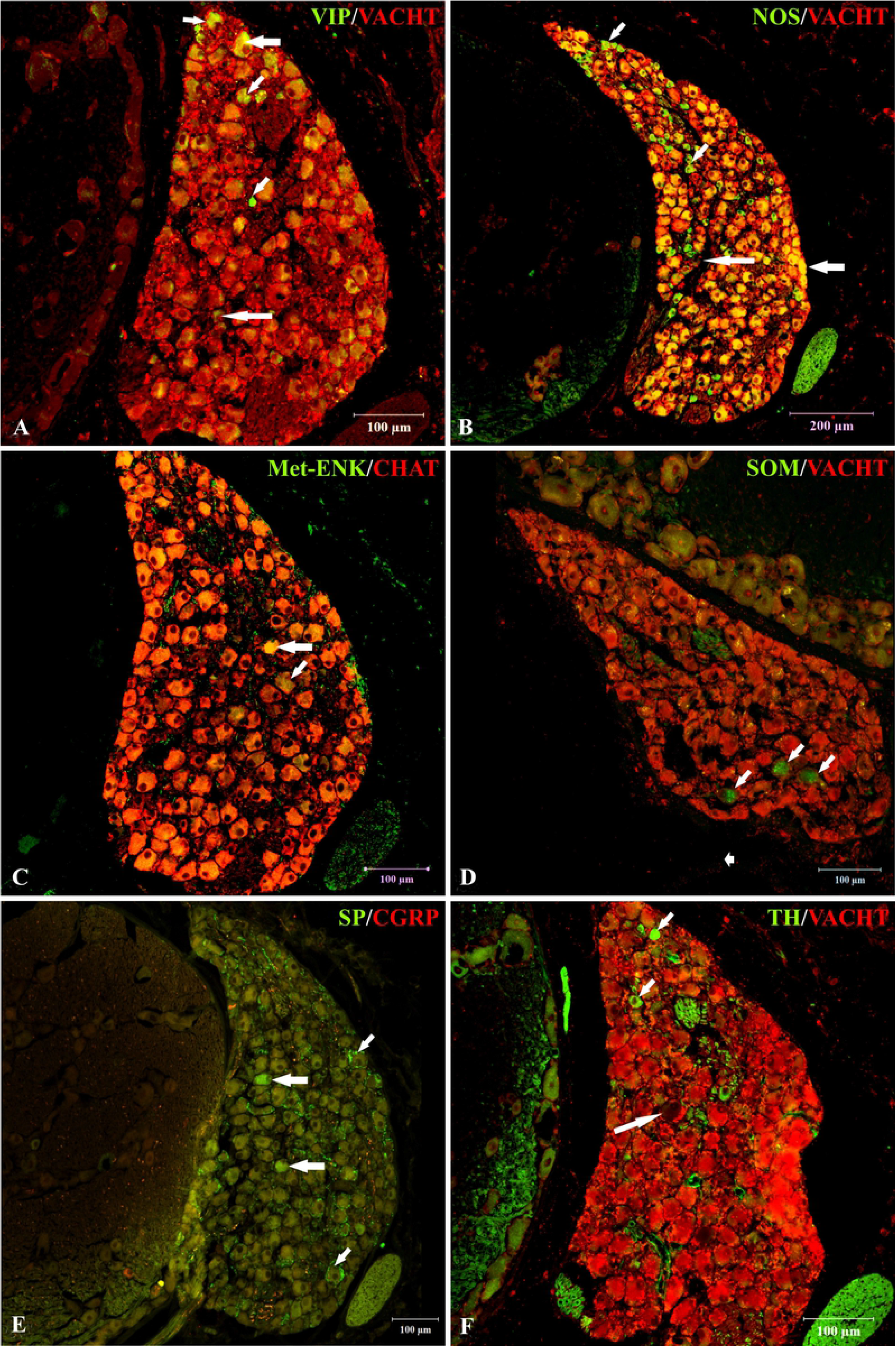
Immunohistochemical coding of the otic ganglion in chinchilla. A) Section from the otic ganglion stained with antibodies against VAChT (red) and VIP (green). The majority of the neurons exhibited immunoreactivity to VAChT, and a number of them also stained to VIP. Some neurons displayed immunoreactivity to VIP only (examples indicated by small arrows). Vast majority of VIP-IR neurons is also VAChT-IR (example indicated by large arrow). Single neurons are immunonegative for both studied substances (long arrow). Very numerous VAChT-immunoreactive nerve fibers formed “basket-like structures” surrounding perikarya. B) Section from the otic ganglion stained with antibodies against VAChT (red) and NOS (green). Over half of the neurons contained NOS-immunoreactivity, whereas double stained nerve cell bodies comprised approx. 45% of all the neurons (examples indicated by large arrow). Neurons stained for NOS only are indicated by small arrows. Single neurons are immunonegative for both studied substances (long arrow). C) ChAT- (red) and Met-ENK-positive (green) nerve structures in the otic ganglion. The vast majority of ganglionic cells were ChAT-positive. Single neurons contained Met-ENK-immunoreactivity (arrows) and some neuronal somata were simultaneously ChAT-positive (large arrow). Part of neurons contained only Met-ENK-IR (small arrow). Note the presence of Met-ENK-IR ganglionic nerve fibers. D) VAChT- (red) or SOM-positive (green) neurons in the otic ganglion. SOM-immunoreactivity was restricted to single perikarya (small arrows) and fine nerve fibers. E) CGRP- (red) and/or SP-positive (green) structures in the otic ganglion. Immunoreactivity to SP was present in single neurons (large arrows). Note the presence of moderate in number SP-positive (green) and /or CGRP-positive (yellow/red) nerve fibers. F) VAChT- (red) or TH-positive (green) neurons in the otic ganglion. Immunoreactivity to TH was present in single perikarya (small arrows). Single neurons are immunonegative for both studied substances (long arrow).

Immunoreactivity to enkephalin was expressed in the nerve cell bodies and nerve fibers. Staining against Met-ENK revealed only 3–4 nerve cell bodies per section that were immunoreactive to or this peptide (Fig 2C); approximately half of these were simultaneously ChAT-positive (Fig 2C). Intraganglionic Met-ENK-IR nerve fibers were moderate in number (Fig 2C). Leu-ENK-immunoreactivity was found in the cell bodies of one or two neurons per section. These perikarya were simultaneously VAChT-positive. The nerve fibers were very scarce.

GAL-IR neurons also occurred only individually (3–4 per section) and stained for ChAT. GAL-positive nerves were very scarce and delicate. Few SOM-positive only nerve cell bodies were encountered in the ganglion (Fig 2D). Intraganglionic SOM-IR nerves were solitary and delicate. SP-positive intraganglionic nerve fibers were numerous and some of them also stained for CGRP (Fig 2E). SP-IR nerves formed basket-like structures surrounding nerve cell bodies. CGRP-immunoreactivity was found in nerve fibers which were less numerous than those which stained for SP. The vast majority of these were also SP-positive; individual nerve fibers contained only CGRP-immunoreactivity. Immunoreactivity to SP was also found in 1–2 neurons per section (Fig 2E). Also, 3–4 neurons per section stained for TH; these were small, intensely stained and scattered throughout the ganglion (Fig 2F).

Our results did not reveal any sex differences.

## Discussion

Our analysis of the topography of the otic ganglion in mammals revealed a close relationship between two anatomical structures: the mandibular nerve and the maxillary artery [14, 3]. In most small mammalian species that have been investigated, the maxillary artery runs along the medial size of the mandibular nerve. In this case, the otic ganglion was located on the medial surface of the mandibular nerve or had direct contact with the maxillary artery [15, 3, 16–18]. The otic ganglion in the chinchilla is an intracranial structure found on the medial surface of the mandibular nerve, and only postganglionic fibers run parallel to the maxillary artery.

In most mammals, the otic ganglion is a single compact aggregation of neurons. This arrangement has been observed in small mammals, including rabbit [19], guinea pig, dog, hedgehog [14, 3, 16,], mouse, rat, hamster, midday gerbil [17, 18], spotted suslik [5], Egyptian spiny mouse [6], and larger mammals [15, 20, 21].

Sometimes the otic ganglion forms a ring-shaped structure encircling the maxillary artery [18]. In some species, the otic ganglion consists of several interconnected nerve cell clusters. This morphological form has been described in the wild pig [22], dog [3], goat, and sheep [23], whereas in cats the ganglion is composed of one main cluster of neurons and about ten small additional ganglia [4]. The otic ganglion in the chinchilla represents a single agglomeration of neurons and only one small additional ganglion was found, associated with the small petrosal nerve.

This relatively small ganglion is an important source of nerve fibers contributing to the innervation of many organs, including the parotid gland, buccal glands, and the blood vessels supplying the brain, lower lip, and gingiva [24, 25]. Previous studies have shown that the vast majority of neurons in the otic ganglion are cholinergic in nature [12]. Correspondingly in the chinchilla, 85% of these neurons are immunoreactive to ChAT or VAChT. The presence of catecholaminergic (TH-positive) neurons within this parasympathetic ganglion may be considered somewhat surprising. This small neuronal population probably represents a kind of the “developmental relic”, consisting of not fully functional adrenergic nerve cells. This has been suggested in relation to catecholaminergic neurons in the porcine ciliary and pterygopalatine ganglia [26, 27], and in the pterygopalatine ganglion of the chinchilla [28]. However, such observations concerning different parasympathetic ganglia have previously also been made regarding pig [26, 12, 27] and rat [8,29,30], as well as human, monkey, guinea pig, ferret, dog, and cat [for references see: 26]. Catecholaminergic neurons usually constitute small subpopulations in these ganglia, but they are much more numerous in human [31] and monkey [30] ciliary ganglia (23% and 68% of ganglionic neurons, respectively).

Our study showed that VIP-positive neurons constituted 10% of all ganglionic neurons. Few VIP-immunoreactive neurons have been found in the porcine and rat otic ganglion [12, 8], while in the human otic ganglion over 90% of neurons are VIP-positive [32]. In the chinchilla pterygopalatine ganglion, a large population (40%) of VIP-immunoreactive neurons was found [28], whereas in the pig, all neurons in this ganglion were VIP-positive [27]. Our study found that many (60%) ganglionic neurons stained for NOS. A large (80%) population of NOS-positive neurons have been found in the human otic ganglion [32]. The pterygopalatine ganglion in the chinchilla also makes up a large population (42%) of NOS-IR neurons [28], whereas all the neurons are NOS-positive in the porcine pterygopalatine ganglion [27].

This study has revealed small populations of neurons immunoreactive to Met-enkephalin and Leu-enkephalin. The expression of opioid peptides in otic ganglion neurons has been studied in the guinea pig [33, 34] and, unlike our results on the chinchilla, a large (35%) subpopulation of the opioidergic neurons was found [34]. Individual GAL- and SOM-IR neurons were found in the porcine otic ganglion [12]. These findings correspond with our results.

In our study, we observed a dense network of VAChT-immunoreactive nerve terminals, frequently forming basket-like structures surrounding the nerve cell bodies. The nerve fibers that were immunoreactive to SP and CGRP were also relatively numerous. It can be assumed that the VAChT-positive nerve fibers represent two populations of nerve terminals: One consists of postganglionic nerve fibers originating from the ganglion, and the second consists of preganglionic nerves originating from the parasympathetic nucleus of the glossopharyngeal nerve. SP and/or CGRP-positive nerve fibers are probably sensory in nature and most likely represent collaterals of dendrites from the mandibular nerve which pass through the otic ganglion.

A comparison of previous findings with our present results suggests the existence of profound interspecies differences in the immunohistochemical characteristics of nerve structures of the otic ganglion. Further studies should be undertaken to clarify the functional significance of these differences.

## Acknowledgements

The authors would like to thank Ms M. Marczak for her excellent technical assistance.

## References

1. Karnovsky MJ, Roots L. A “direct-coloring” thiocholine method for cholinesterases. J Histochem Cytochem. 1964;12: 219–221.

2. Gienc J. The application of the histochemical method in the anatomical studies on the parasympathetic ganglia and nerve bundles of postganglionic axons in the sublingual region of some mammals. Zool Pol. 1977;26: 187–192.

3. Gienc J. Kuder T. Otic ganglion in dog. Topography and macroscopic structure. Folia Morphol (Warsz.). 1983;42: 31–40.

4. Kuchiiwa S, Kuchiiwa T, Nonaka S, Nakagawa S. Facial nerve parasympathetic preganglionic afferents to the accessory otic ganglia by way of the chorda tympani nerve in the cat. Anat Embryol (Berl.). 1998;197: 377–382.

5. Szczurkowski A. Morphology, topography and cytoarchitectonics of the otic ganglion in the spotted suslik (Spermophilus suslicus, Guldenstaedt 1770). Ann Anat. 1999;181: 409–411.

6. Szczurkowski A, Kuder T, Nowak E, Kuchinka J. Morphology, topography and cytoarchitectonics of the otic ganglion in Egyptian spiny mouse (Acomys cahirinus, Desmarest). Folia Morphol. (Warsz.). 2001;60: 61–64.

7. Shimizu T. Distribution and pathway of the cerebrovascular nerve fibers from the otic ganglion in the rat: anterograde tracing study. J Auton Ner. Syst. 1994;49: 47–54.

8. Hardebo JE, Suzuki N, Ekblad E, Owman C. Vasoactive intestinal polypeptide and acetylcholine coexist with neuropeptide Y, dopamine-beta-hydroxylase, tyrosine hydroxylase, substance P or calcitonin gene-related peptide in neuronal subpopulations in cranial parasympathetic ganglia of rat. Cell Tissue Res. 1992;267: 291–300.

9. Leblanc GG, Trimmer BA, Landis SC. Neuropeptide Y-like immunoreactivity in rat cranial parasympathetic neurons: coexistence with vasoactive intestinal peptide and choline acetyltransferase. Proc Natl Acad Sci. 1987;84: 3511–3515.

10. Suzuki N, Hardebo JE, Kahrstrom J, Owman C. Neuropeptide Y co-exists with vasoactive intestinal polypeptide and acetylcholine in parasympathetic cerebrovascular nerves originating in the sphenopalatine, otic, and internal carotid ganglia of the rat. Neuroscience. 1990a;36: 507–519.

11. Suzuki N, Hardebo JE, Owman C. Origins and pathways of choline acetyltransferase-positive parasympathetic nerve fibers to cerebral vessels in rat. J Cereb Blood Flow Metab. 1990b;10: 399–408.

12. Lakomy M, Kaleczyc J, Wasowicz K, Czaja K. Immunohistochemical study of the otic ganglion in the pig. Pol J Vet Sci. 2002;5: 257–262.

13. Ayer-LeLievre C, Seiger A. Substance P-like immunoreactivity in developing cranial parasymphatetic neurones of the rat. Int J Devl Neuroscience. 1985;3: 267–277.

14. Gienc J, Kuder T. Morphology and topography of the otic ganglion in guinea pig detected with thiocholine technique. Folia Morphol (Warsz.). 1980;39: 79–85.

15. Fisbach I, Dudzińska B. Topografia zwoju usznego u królika. Folia Morphol (Warsz.). 1970;21: 241–247.

16. Gienc J, Kuder T, Szczurkowski A. Parasympathetic ganglia in the head of western hedgehog (Erinaceus europaeus). I. Otic ganglion. Acta Theriol. 1988;33: 115–120.

17. Kuder T. Comparative morphology and topography of cranial parasympathetic ganglia connected with the trigeminal nerve in mouse, rat and hamster (Mus musculus L. 1759, Rattus norvegicus B. 1769, Mesocricetus aureatus W. 1839). Part I. Otic ganglion. Folia Morphol (Warsz.). 1983;42: 187–197.

18. Kuder T. Topography and macroscopic structure of parasympathetic ganglia connected with the trigeminal nerve in midday gerbil (Meriones meridianus - Mammalia: Rodentia). Acta Biol Cracow Zool. 1985;27: 61–71.

19. Dixon JS. The fine structure of parasympathetic nerve cells in the otic ganglia of the rabbit. Anat Rec. 1966;156: 239–252.

20. Godinho HP. A comparative anatomical study of cranial nerves in goat, sheep and bovine (*Capra hirus*, *Ovis aries* and *Bos taurus*) their distribution and autonomic components. Iowa State University, Ames-Iowa. 1968.

21. Petela L. Topografia nerwu trójdzielnego u bydła. Część III. Nerw żuchwowy. Pol Arch Vet. 1974;17: 559–580.

22. Petela L. Nerw trójdzielny u dzika (*Sus scrofa* L. 1758). Zesz Nauk AR Wrocław. 1979;17: 1–55.

23. Kovšikova LP. Usnoj uzel - gln. oticum domasnich żivotnych. Ucennyje Zap Viteb Vet Inst. 1958;16: 11–114.

24. Edvinsson L, Elsas T, Suzuki N, Shimizu T, Lee TJ. Origin and Co-localization of nitric oxide synthase, CGRP, PACAP, and VIP in the cerebral circulation of the rat. Microsc Res Tech. 2001;53: 221–228.

25. Shimizu T, Koto A, Suzuki N, Morita Y, Takao M, Otomo S, et al. Occurrence and distribution of substance P receptors in the cerebral blood vessels of the rat. Brain Res. 1999;830: 372–378.

26. Kaleczyc J, Juranek J, Calka J, Lakomy M. Immunohistochemical characterization of neurons in the porcine ciliary ganglion. Pol J Vet Sci. 2005;8: 65–72.

27. Podlasz P, Wąsowicz K, Kaleczyc J, Lakomy M, Bukowski R. Localization of immunoreactivities for neuropeptides and neurotransmitter-synthesizing enzymes in the pterygopalatine ganglion of the pig. Vet Med – Czech. 2003;48: 99–107.

28. Szczurkowski A, Sienkiewicz W, Kuchinka J, Kaleczyc J. Morphology and immunohistochemical characteristics of the pterygopalatine ganglion in the chinchilla (Chinchilla laniger, Molina). Pol J Vet Sci. 2013;16: 359–368.

29. Landis SC, Jackson PC, Fredieu JR, Thibault J. Catecholaminergic properties of cholinergic neurons and synapses in adult rat ciliary ganglion. J Neurosci. 1987;7: 3574–3587.

30. Uemura Y, Sugimoto T, Nomura S, Nagatsu I, Mizuno N. Tyrosine hydroxylase-like immunoreactivity and catecholamine fluorescence in ciliary ganglion neurons Brain Res. 1987;416: 200–203.

31. Kirch W, Neuhuber W, Tamm ER. Immunohistochemical localization of neuropeptides in the human ciliary ganglion. Brain Res. 1995;681: 229–234.

32. Uddman R, Tajti J, Moller S, Sundler F, Edvinsson L. Neuronal messengers and peptide receptors in the human sphenopalatine and otic ganglia. Brain Res. 1999;826: 193–199.

33. Gibbins IL. Target-related patterns of co-existence of neuropeptide Y, vasoactive intestinal peptide, enkephalin and substance P in cranial parasympathetic neurons innervating the facial skin and exocrine glands of guinea-pigs. Neuroscience. 1990;38: 541–560.

34. Shimizu T, Morris JL, Gibbins IL. Expression of immunoreactivity to neurokinin-1 receptor by subsets of cranial parasympathetic neurons: correlation with neuropeptides, nitric oxide synthase, and pathways. Exp Neurol. 2001;172: 293–306.

